# PARP1 deficiency induces aging-associated cardiac failure via activation of Akt signalling

**DOI:** 10.64898/2025.12.01.691751

**Authors:** Thoniparambil Sunil Sumi, Sneha Mishra, Amarjeet Shrama, Abhishek Chowdhury, Bhoomika Shivanaiah, Dimple Nagesh, Souvik Roy, Aastha Munjal, Ashwini Yogilal Bhelave, Kumaravel Somasundaram, Nagalingam Ravi Sundaresan

**Author notes:** Corresponding author: Mailing address: Nagalingam R. Sundaresan, Associate Professor, Lab SB-02, Department of Microbiology and Cell Biology, Division of Biological Sciences, Indian Institute of Science, Bengaluru – 560012, Karnataka, India. Contributed equally as first authors (Thoniparambil Sunil Sumi, Sneha Mishra).

## Abstract

PARP1, a poly-ADP-ribose transferase, plays a critical role in maintaining genomic stability, transcription, cellular metabolism, and cell death. PARP1 inhibition protects cardiomyocytes against oxidative and genotoxic stress. However, the role of PARP1 in aging-associated heart failure remains poorly explored. In the current study, we report that PARP1 levels are downregulated in aging mouse hearts, and PARP1 deficiency induces aging-related cardiac remodelling and contractile dysfunction in mice. PARP1 deficient mice hearts exhibit spontaneous activation of the Akt signalling pathway, leading to the development of aging-related cardiac hypertrophy and fibrosis. Our findings reveal two distinct mechanisms by which PARP1 regulates Akt signalling, direct interaction of PARP1 with Akt and the transcriptional regulation of phosphatases like PTEN, a negative regulator of Akt signalling. PARP1 binds and inhibits Akt by poly-ADP-ribosylation at E40 and E49 residues, which impairs Akt membrane recruitment and subsequent activation. Inhibition of Akt reversed hypertrophy in PARP1-depleted cardiomyocytes and improved the contractile dysfunction in PARP1-deficient hearts. These findings reveal a previously unrecognized regulatory role for PARP1 in aging-associated cardiac failure.

## Introduction

Cardiovascular diseases are becoming increasingly common with an expanding elderly population. During aging, the heart undergoes intrinsic modifications that weaken its normal function [1]. With advancing age, left ventricular hypertrophy becomes increasingly common, evidenced by progressive thickening of the ventricular wall. This age-related remodelling compromises cardiac performance, leading to elevated morbidity and mortality among the elderly [2]. Even though the medical field has made a huge progression in the management of the risk factors associated with cardiovascular disease, there has been no cure for cardiovascular diseases, and it still remains the leading cause of mortality, morbidity, and disability [3].

NAD⁺-dependent enzymes play a central role in regulating cardiac function [4, 5]. Poly (ADP-ribosylation (PARylation) is an NAD⁺-dependent post-translational modification in which multiple ADP-ribose units are transferred to target proteins. This reaction is primarily catalysed by members of the PARP enzyme family [6, 7]. PARP1, a major Poly(ADP-ribose) transferase enzyme, is localized in the nucleus and is involved in maintaining genomic stability, transcription, cellular energy metabolism, and cell death [8]. PARP1 inhibition has been previously reported to protect against multiple cardiac pathologies, including cardiac reperfusion injury and cardiac hypertrophy [9]. In contrast, PARP1 knock-out mice exhibit an increased incidence of tumors, accelerated aging, and a reduced lifespan [10]. Even though previous studies suggest that PARP1 inhibition is beneficial against oxidative stress and protects against starvation, the role of PARP1 in aging-associated heart failure remains unexplored.

The molecular basis of cardiac aging has been linked to the dysregulation of various key signalling networks, including the insulin/IGF–PI3K pathway, adrenergic signalling, the renin– angiotensin system, and nutrient-responsive signalling cascades [2]. Alterations in the activity of Akt kinase (also known as protein kinase B) have been implicated in aging and aging-related diseases [11–13]. Akt is an integral part of the insulin/IGF signalling pathway, which regulates cellular homeostasis [14]. Experimental evidence suggests that the Akt1 isoform is necessary for growth, as mice deficient in Akt1 show both foetal and postnatal growth defects [15]. Additionally, numerous studies have demonstrated the essential role of Akt in cell survival, growth, and proliferation [14, 16, 17]. Akt has been extensively studied due to its ability to transform cells, and Akt gene amplification is also a key event in several tumours [18, 19]. One of the well-studied nuclear substrates of Akt is the FoxO (forkhead box protein <u>O</u>) family of transcription factors. Akt phosphorylates FoxO inside the nucleus, promotes its nuclear export to the cytoplasm, where it gets degraded [20–22]. Impaired nuclear Akt activity leads to chronic hyperactivation of FoxO in the nucleus [23, 24]. Additionally, it phosphorylates the GSK-3β kinase at Serine 9, thereby inhibiting its catalytic activity [16, 25].

Under basal conditions, Akt is localized in the cytoplasm and in the nucleus, where post-translational modifications of Akt, especially phosphorylation, ubiquitination, and acetylation, play a key role in the activation of Akt [26–31]. Phosphorylation of Akt at Thr^308^ and Ser^473^ residues is absolutely essential for its kinase activity [27]. Phosphorylation and activation of Akt is a multistep process, which involves (a) membrane recruitment of Akt, (b) conformational change at the membrane, (c) phosphorylation by upstream kinases, and (d) translocation to cytoplasm and nucleus [27, 32]. Our previous work identified acetylation as a critical regulator of PIP_3_ binding to the N-terminal Pleckstrin homology (PH) domain and membrane recruitment of Akt [28]. Another mechanism established for AKT membrane recruitment is by ubiquitination through TRAF6, E3 ubiquitin ligase [29]. Our recent data demonstrate that PARP1 modulates Akt through PARylation, thereby reducing membrane localization and activation of Akt [33]. Inhibition of PARP1, therefore, augments Akt signalling and confers protection against JEV [33]. However, the role of PARP1 in regulating Akt and thereby modulating the aging-associated cardiac remodelling and failure has not been well explored yet.

In the present study, we investigated the role of PARP1 in regulating cardiac aging using mouse models and aimed to characterize the Akt residues that are regulated by PARP1. Our findings suggest that PARP1 deficiency spontaneously activates the Akt signalling pathway, leading to the development of aging-related cardiac failure. PARP1 negatively regulates Akt activation by PARylating Akt at E40 and E49 residues, thereby suppressing its membrane localization.

## Results

### PARP1 deficiency induces spontaneous age-dependent cardiac remodelling and failure

Several studies demonstrate that with age, cardiomyocytes undergo hypertrophic changes concomitant with an increase in the proliferation of cardiac fibroblasts, ultimately causing heart failure [2, 34]. To determine if PARP1 levels are affected in the hearts of aged mice, we examined PARP1 protein levels in the hearts of mice from different age groups. Our findings suggest that PARP1 protein levels are significantly downregulated in the hearts of aged mice **(Figures 1A, B)** and correlate with the age-associated cardiac dysfunction in mice [35]. We hypothesized that depletion of PARP1 may contribute to the development of cardiac hypertrophy and heart failure in aged mice. To verify this, we studied the whole-body PARP1 knockout mice (PARP1-KO) **(Figure 1C)** and assessed their cardiac function in an age-dependent manner using echocardiography. At two months of age, the heart weight-to-tibia length ratio of PARP1-KO mice was also comparable to that of the wild-type mice **(Figure 1D)**. We did not observe any changes in cardiac function in two-month-old young PARP1-KO mice, as assessed by echocardiography **(Figures 1E-H)**. Additionally, we examined cardiomyocyte size in the hearts of young PARP1-KO mice and observed that it was comparable to that of WT mice **(Figures 1I-J)**, suggesting that the hearts of PARP1-KO mice function normally at a young age. However, one-year-old PARP1-KO mice spontaneously developed adverse cardiac remodelling with enlarged hearts and an increased heart-to-tibia length ratio **(Figures 1K, L)**. Echocardiography analysis revealed a reduction in ejection fraction, fractional shortening, and left ventricular posterior wall thickness, accompanied by an increase in left ventricular internal diameter (**Figures 1M-P**), indicating the development of cardiomyopathy with contractile dysfunction. We further observed signs of fibrotic tissue deposition in heart sections **(Figures 1Q, R)**. These one-year-old PARP1-KO hearts also exhibited an increase in cross-sectional area, indicating cardiomyocyte hypertrophy **(Figure 1S, T)**. Together, these results suggest that PARP1 deficiency results in spontaneous cardiac hypertrophy and heart failure in an age-dependent manner.

**Figure 1:**
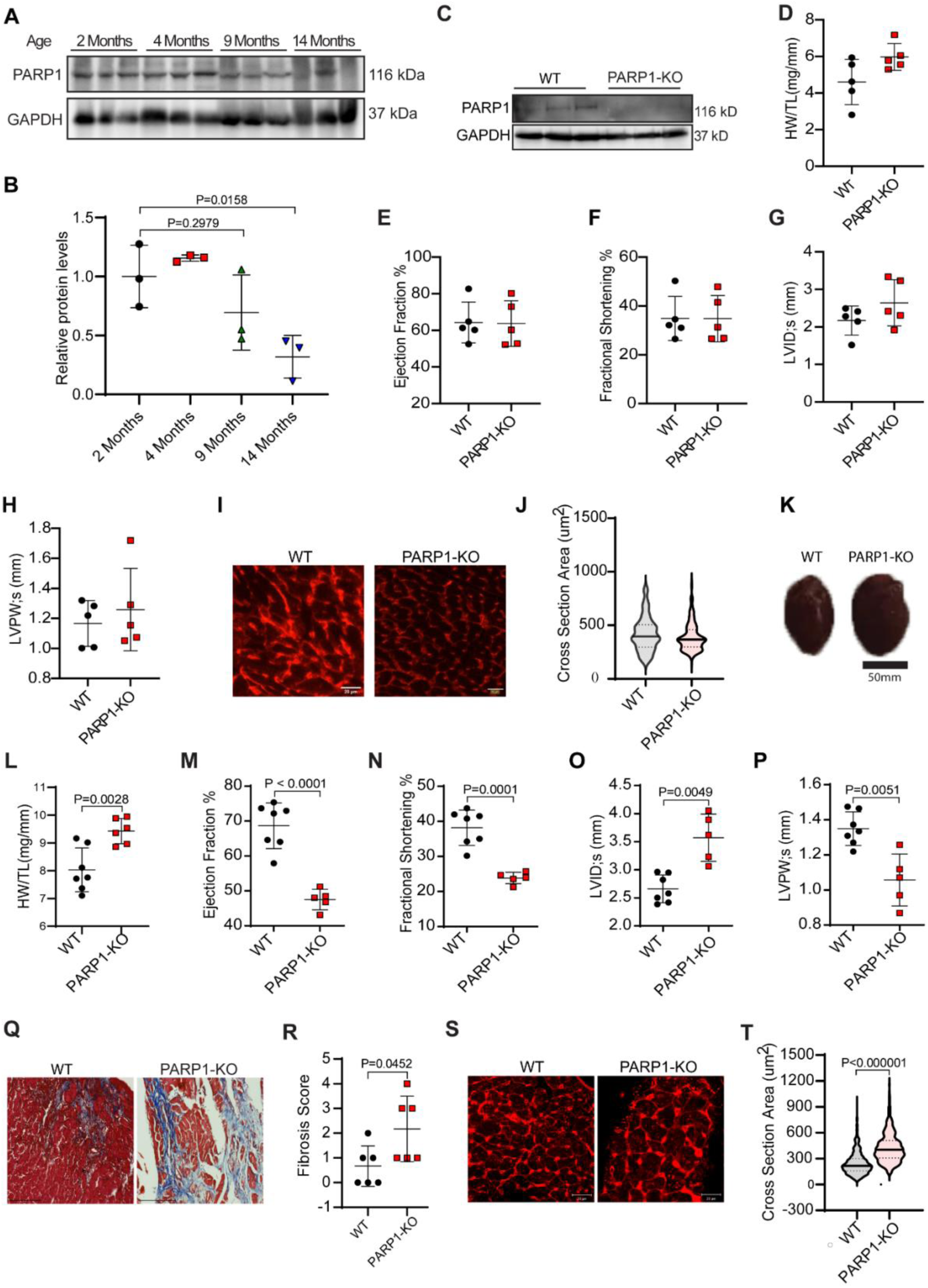
PARP1-KO mice hearts exhibit cardiac hypertrophy. **(A)** Representative immunoblot showing PARP1 expression in heart tissue of different age groups, 2, 4, 9, and 14 months of WT or PARP1-KO mice. n=3 mice per group. **(B)** Quantitative representation of PARP1 levels in hearts of different-aged mice depicted in Figure 1A. **(C)** Representative immunoblot showing PARP1 expression in heart tissue of WT or PARP1-KO mice. Indicated sizes are in kilodaltons (kDa), n=3 mice per group. **(D)** Scatter plot representing the HW/TL ratio of WT and PARP1-KO mice at 2 months of age, n = 6-7 mice per group. (**E-H)** Scatter plots depicting **(E)** Ejection Fraction **(F)** Fractional Shortening **(G)** Left Ventricular Internal Diameter systole, LVID; s **(H)** Left Ventricular posterior Wall Thickness, LVPW; s of WT and PARP1-KO mice at 2 months of age, n = 5 mice per group. **(I)** Representative confocal images of WGA-stained cardiac tissue sections showing the cross-sectional area of cardiac cells in WT and PARP1-KO mice at 2 months of age. *Scale bar* = 20 μm. **(J)** Violin plot showing quantification of WGA images represented in Figure 1(I). n= 111(Control), 248(PARP1-KO), cells per group. **(K)** Representative image of heart isolated from WT and PARP1-KO mice of 14 months age, showing hypertrophic growth in the heart of PARP1-KO mice when compared with control (WT) mice. **(L)** Scatter plot representing HW/TL ratio of WT and PARP1-KO mice at 14 months of age, n = 6-7 mice per group. **(M-P**) Scatter plots depicting (**M**) Ejection Fraction (**N**) Fractional Shortening (**O**) Left Ventricular Internal Diameter systole, LVID;s **(P)** Left Ventricular posterior Wall Thickness, LVPW;s of WT and PARP1-KO mice at 14 months of age, n = 5-7 mice per group. **(Q**) Representative Fluorescence microscopic images of Masson’s Trichrome Stained heart tissue sections of WT and PARP1-KO mice of 3 months of age, n=6. *Scale bar* = 150 μm. **(R)** Quantification of Masson’s Trichrome images represented in Figure 1 (Q), showing fibrosis score in cardiac tissue of WT and PARP1-KO mice of 14 months of age. Scoring was done blindly. Data are shown as mean ± s.d. n=6 mice per group**. (S)** Representative confocal images of WGA-stained cardiac tissue sections showing the cross-sectional area of cardiac cells in WT and PARP1-KO mice at 14 months of age. *Scale bar* = 20 μm. **(T)** Violin plot showing quantification of WGA images represented in Figure 1(s). n= 732(Control), 502(PARP1-KO), cells per group Data information: In **(B)**, P values are shown from Ordinary one-way ANOVA with Dunnett’s multiple comparisons test, data are represented as mean ± S.D.*P≤0.05, n=3 mice per group. **(D-H)** data are represented as mean ± S.D.*P≤0.05; Student’s t-test with Welch correction was used to calculate the p-values, n=5 animals per group. In **(L-P, R)** data are represented as mean ± S.D.*P≤0.05; Student’s t-test with Welch correction was used to calculate the p-values, n=5-7 animals per group. In (**J and T**) P values shown are from an unpaired, two-tailed t-test (Mann-Whitney test). Data are presented as medians with 25th and 75th percentiles. N=5 mice per group, n= 111(Control), 248 (PARP1-KO) for two months and n= 732(Control), 502(PARP1-KO) for fourteen months cells per group.

### PARP1 deficiency spontaneously hyperactivates Akt signalling in the heart

We performed RNA sequencing on heart tissues from control and PARP1-KO mice to explore the underlying molecular mechanisms involved in spontaneous cardiac hypertrophy **(Figures 2A-D).** Several pathways associated with cardiac hypertrophy were significantly deregulated in PARP1-deficient hearts. Notably, pathways associated with the negative regulation of PI3K-Akt network, insulin receptor signalling, and metabolism were significantly downregulated, while calcium regulation in cardiac cells and Notch signalling pathways were significantly upregulated **(Figure 2D).** It is well-established that Akt signalling is critical in regulating cardiac pathophysiology [36]. Thus, to further investigate the molecular mechanism via which PARP1 deficiency may perturb cardiomyocyte homeostasis and induce cardiac hypertrophy, we first evaluated the expression of phospho-Akt (S473) and phospho-Akt (T308) as markers of Akt activation in the heart tissues derived from wild-type control and PARP1-KO mice. The expression of both phospho-Akt (S473) and phospho-Akt (T308) was upregulated in the hearts of PARP1-KO mice, indicating Akt activation **(Figures 3A, B)**. Activated Akt is known to phosphorylate and regulate its downstream targets, FoxO1 and GSK-3β. Thus, to verify Akt activation under PARP1-KO conditions, we assessed the phosphorylation of these targets in control and PARP1-KO hearts and found increased phosphorylation of these targets in PARP1-KO hearts **(Figures 3A, B)**, indicating that Akt is indeed activated upon PARP1 depletion.

**Figure 2:**
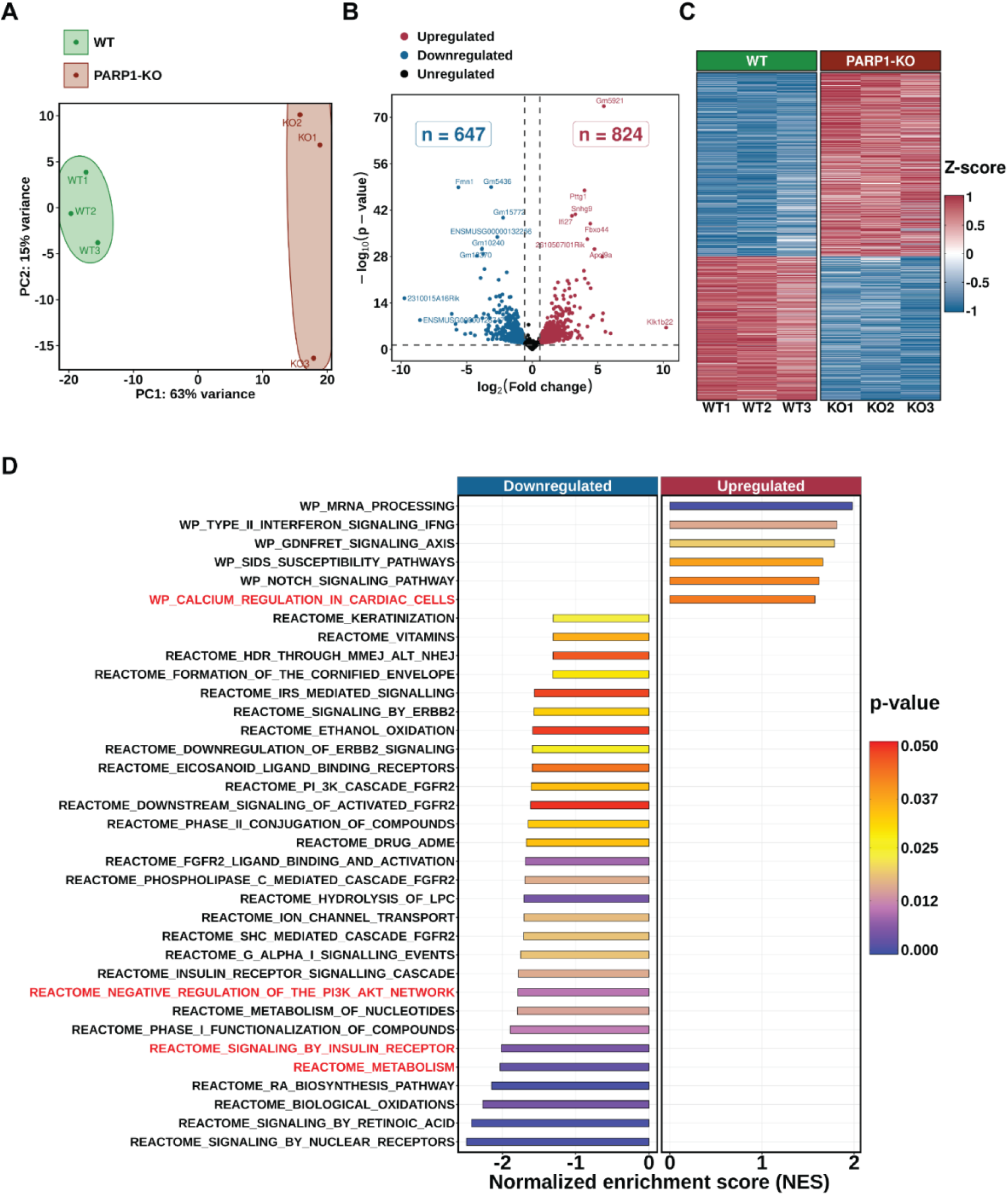
Akt signalling pathway genes are upregulated in mice hearts lacking PARP1. **(A)** Principal Component Analysis (PCA) of gene expression data from wild-type (WT) and PARP1-KO mice hearts, with three replicates per condition. Points represent individual replicates, and colours indicate the two experimental conditions. The scatter plot demonstrates separation of the two groups, with the first two principal components (PC1 and PC2) accounting for the primary variance in the data. **(B)** Volcano plot illustrating differential gene expression between WT and PARP1-KO groups. The cutoffs for upregulation and downregulation are set at log_2_ fold changes of 0.58 and −0.58, respectively. Significantly regulated genes (p-value < 0.05) with large fold changes are labelled. **(C)** Heatmap showing the Z-score normalized gene expression in WT and PARP1-KO mice hearts, each with three biological replicates. The Z-scores are calculated for each gene, and the colour scale represents relative expression, with red indicating upregulation and blue indicating downregulation. **(D)** Bar diagram illustrating the upregulated (right) and downregulated (left) pathways identified through Gene Set Enrichment Analysis (GSEA). Pathways are arranged from top to bottom in descending order of their normalized enrichment score (NES). Bar colour reflects the p-value of the pathway. Pathways of relevance to this study are highlighted in red.

**Figure 3.**
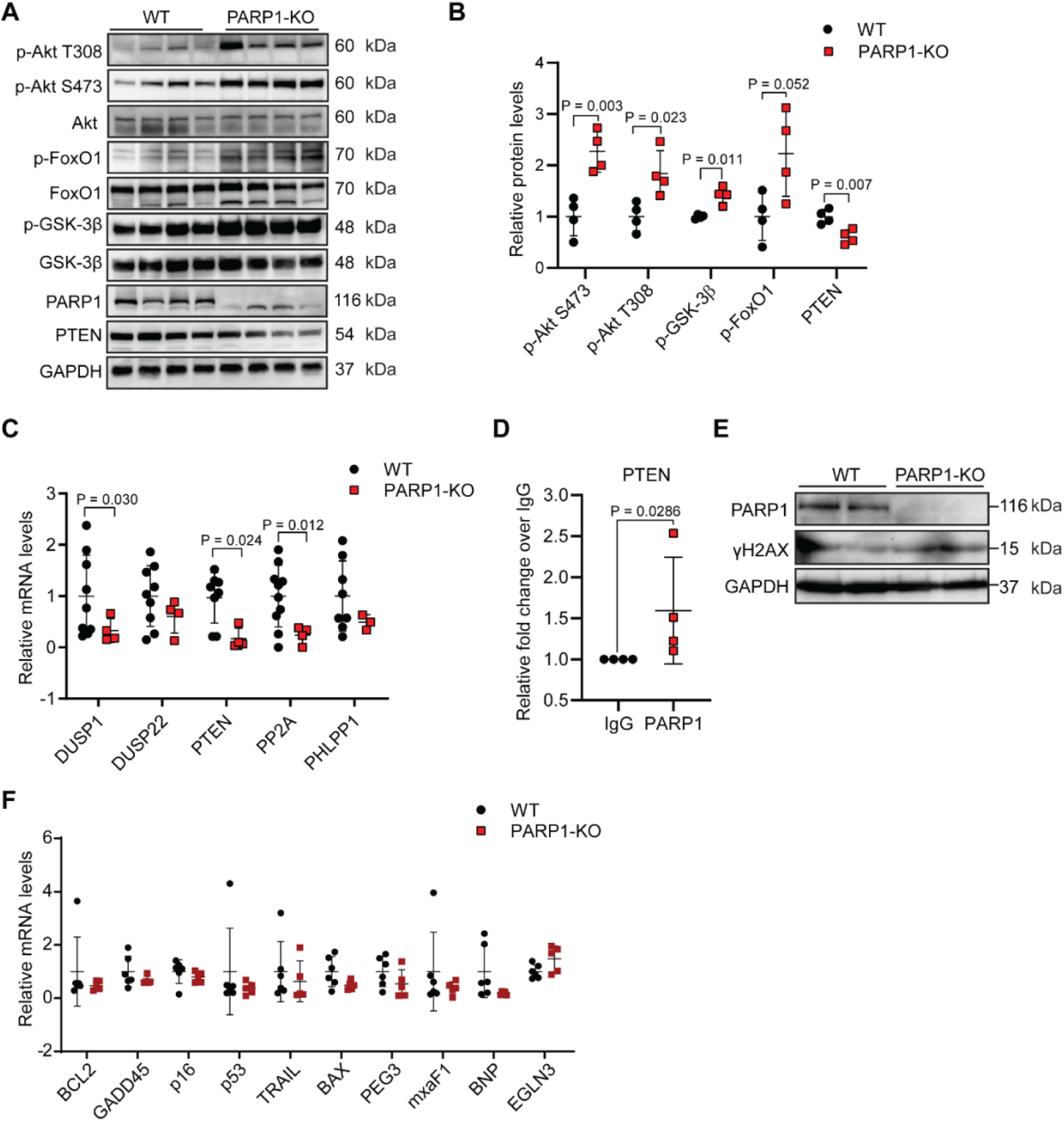
PARP1 deficiency spontaneously hyperactivates Akt signalling in the heart. **(A)** Representative images of immunoblot analysis in heart tissue of WT and PARP1-KO mice showing hyperactivated Akt pathway in PARP1-KO mice when compared with WT mice, n=4 animals per group. (**B)** Quantitative representation of the Akt pathway analysis shown in Figure 3(A). (**C)** qPCR analysis of relative mRNA expression levels of various phosphatases (as indicated in the figure) in WT and PARP1-KO mice heart tissues. n = 4–10 mice per group. (**D)** Quantitative representation of CHIP assay indicating relative PARP1 abundance on PTEN promoter, values are represented as fold change relative to IgG control. n=4 animals per group. (**E)** Representative images of immunoblot analysis in heart tissue of WT and PARP1-KO mice showing expression of γH2AX when compared with WT mice, n=2 animals per group. **(F)** qPCR analysis of relative mRNA expression levels of various cell death markers (as indicated in the figure) in WT and PARP1-KO mice heart tissues. n = 5–6 mice per group. Data information: In **(B)** data are represented as mean ± S.D.*P≤0.05; Student’s t-test with Welch correction was used to calculate the p-values. n=4 animals per group. In **(C)** data are represented as mean ± S.D.*P≤0.05; Student’s t-test with Welch correction was used to calculate the p-values for normally distributed datasets, the Mann-Whitney test was used to calculate p-values when the datasets are not normally distributed (PTEN). n = 4–10 mice per group. In **(D)** data are represented as mean ± S.D.*P≤0.05 The Mann-Whitney test was used to calculate p-values. n=4 animals per group. In **(F)** data are represented as mean ± S.D.*P≤0.05; Student’s t-test with Welch correction was used to calculate the p-values for normally distributed datasets, the Mann-Whitney test was used to calculate p-values when the datasets are not normally distributed. n = 5-6 mice per group.

### PARP1 deficiency downregulates the expression of inhibitory Akt phosphatases

To understand how PARP1 regulates Akt activation in the heart, we evaluated the expression of known Akt regulatory phosphatases, including DUSP1, DUSP22, PTEN, PP2A, and PHLPP1 [37–41]. We observed a significant decrease in the levels of several phosphatases, including DUSP1, PTEN, and PP2A mRNA in the hearts of PARP1-KO mice **(Figure 3C)**. We also observed significant downregulation of PTEN protein levels (Figures 3A, B), indicating that PARP1 may regulate Akt activity by modulating PTEN expression. We performed ChIP analysis in the hearts of mice to evaluate whether PARP1 transcriptionally regulates PTEN. ChIP analysis revealed that PARP1 binds to the PTEN promoter, suggesting that PARP1 regulates Akt, at least in part, by transcriptionally regulating the expression of its inhibitory phosphatase regulator, PTEN **(Figure 3D).** The cardiac dysfunction observed in PARP1-KO hearts could potentially result from the loss of the PARP1-mediated DNA damage response. Thus, to acknowledge this possibility, we assessed the levels of γH2AX, a marker of DNA damage **(Figure 3E)**, and various cell death markers, including BCL2, GADD45, p16, p53, TRAIL, BAX, PEG3, mxaF, BNP, and EGLN3 **(Figure 3F)** in PARP1-KO mice, but observed no significant changes suggesting that the observed effect may not be produced due to cardiomyocyte cell death. Altogether, these results suggest that PARP1 may regulate cardiac aging by modulating Akt signalling in the heart.

### PARP1 binds to and PARylates Akt at E40 and E49 residues

Our recent study has found that PARP1 binds to and PARylates the PH domain of Akt, regulating Akt membrane recruitment and acting as a crucial regulator of Akt signalling [33]. It is possible that this mechanism may also operate in the heart. Our findings suggest that PARP1 interacts with Akt, as assessed by co-immunoprecipitation (**Figure 4A**) and confirmed by confocal microscopy (**Figure 4B**). Next, we tested whether Akt is a PARylated protein in cells and tissues. Our findings suggest that Akt is PARylated in cells as well as in different mouse tissues **(Figures 4C, D)**. Our previous findings suggested that PARP1 PARylates the PH domain of Akt, although the specific residues involved were not characterized. In this work, we performed site-directed mutagenesis to generate several mutants at glutamate residues in the PH domain of Akt1. We mutated glutamate to Alanine (E to A), as glutamate is preferentially PARylated by PARP1. We transiently overexpressed wild-type T7-tagged Akt or T7-tagged Akt mutants and assessed the status of PARylation. Upon screening several Akt mutants, we identified that the residues E40 and E49 are PARylated in Akt, as evidenced by the E40A and E49A mutants of Akt exhibiting reduced PARylation, increased phosphorylation, and activity **(Figure 4E).** Next, we were interested in finding how PARylation regulates Akt activation. As reported in literature, Akt activation requires membrane translocation, conformation change, and subsequent phosphorylation [27]. We assessed the membrane localization of Akt E40A, Akt E49A, Akt E17K, and Akt K14Q mutants in serum-starved as well as serum-stimulated cells **(Figure 4F).** It is well known that Akt E17K is constitutively membrane localised and hyperactive in either serum-starved or serum-induced conditions [28], and Akt K14Q mutant is known to be constitutively inactive even on serum induction, as it does not bind to PIP_3_ [28]. Interestingly, our results suggest that Akt-PH E40A and Akt-PH E49A are constitutively membrane-localized, similar to Akt E17K (Figure 4F), which may be the reason for the increased phosphorylation and activity of Akt E40A and E49A mutants. Collectively, the results suggest that Akt is more recruited to the membrane in PARP1-depleted cells, which might be due to reduced PARylation at the E40 and E49 residues present in the PH domain of Akt.

**Figure 4.**
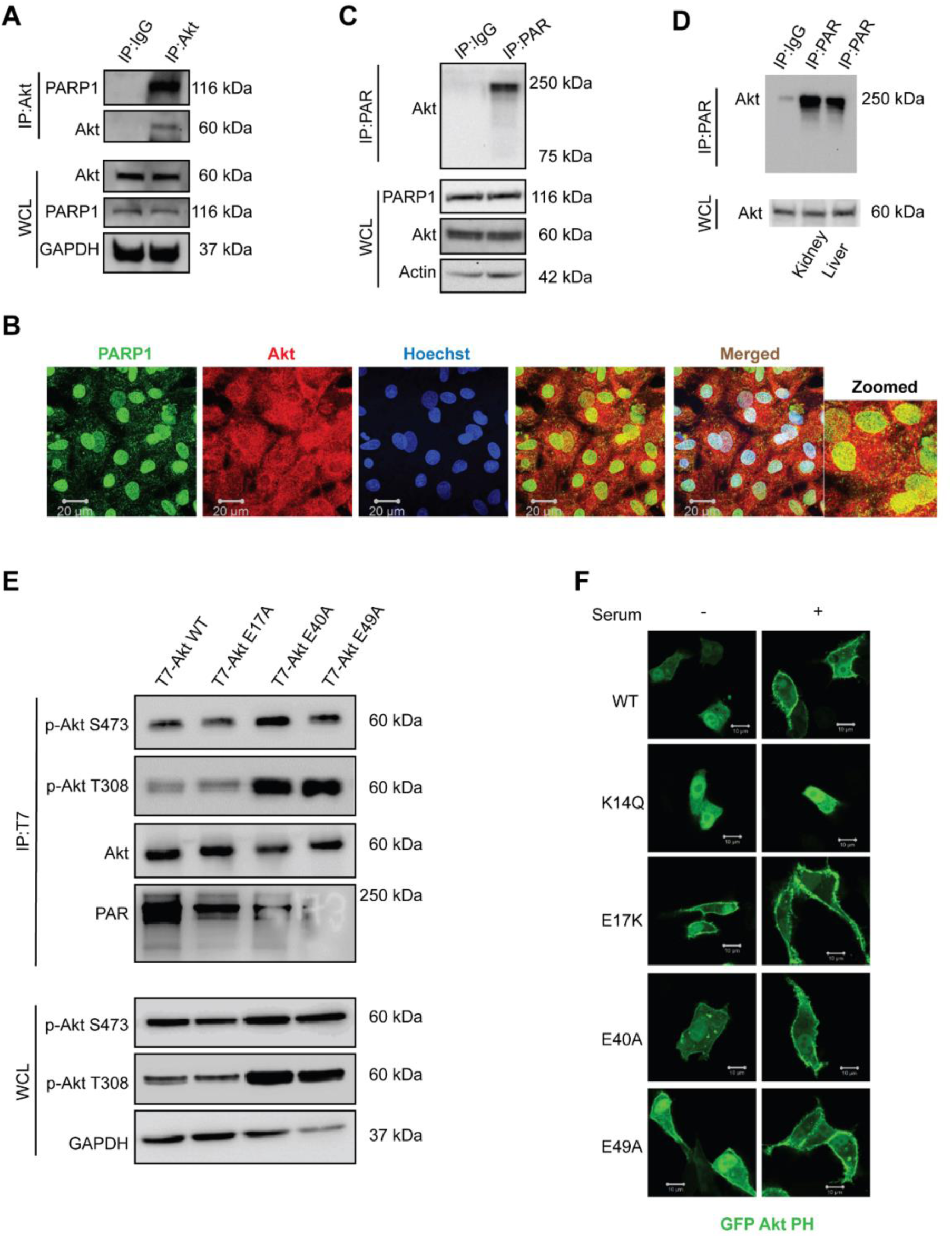
PARP1 binds to and PARylates Akt at E40 and E49 residues. **(A)** Representative immunoblot images showing Akt interaction with PARP1. Akt was immunoprecipitated from 293 cell lysate using Akt specific antibody. Goat-IgG was used as control to eliminate the possibility of non-specific interaction. Bead bound protein and whole cell lysate were subjected to immunoblotting and analysed using Akt, PARP1 antibody. GAPDH was used as loading control. **(B)** Representative immunofluorescence images depicting Akt and PARP1 co-localization inside nucleus. 293 cells were fixed and stained using primary antibodies against Akt and PARP1 and secondary antibodies conjugated to fluorophores were used for confocal microscopy. DAPI was used for nucleus staining. Images were taken at 63X. **(C)** Representative immunoblot images showing Akt as a PARylated protein. Protein lysates of 293 cells were immunoprecipitated for PARylated proteins using PARylation specific antibody and subjected to western blotting for assessing Akt PARylation. **(D)** Akt is a PARylated protein. Protein lysates of mouse kidney and liver tissues were immunoprecipitated for PARylated proteins using PARylation specific antibody and subjected to western blotting for assessing Akt PARylation. **(E)** Representative immunoblot images showing decreased PARylation and increased phosphorylation of Akt in Akt E40A and E49 mutants. 293 cells were transfected with T7 tagged Akt1 WT or either of Akt1 mutants, T7-Akt1 E17A, T7-Akt1 E40A and T7-Akt1 E49A. Akt was immunoprecipitated using T7 specific antibody and subjected to western blotting for assessing Akt PARylation and phosphorylation. **(F)** Representative confocal images showing that Glutamate PARylation in PH domain regulates activation of Akt. 293 cells were transfected with GFP tagged Akt1 PH or GFP Akt1 E17K, GFP Akt1 E40A, GFP Akt1 E49A, GFP Akt1 K14Q. Cells were induced with serum. Confocal images were taken at 63X. GFP Akt1 E17K and GFP Akt1 K14Q were used as positive and negative controls respectively.

### PARP1 depletion induces cardiomyocyte hypertrophy in a cell-autonomous manner

To understand the effect of PARP1 deficiency in terminally differentiated cardiomyocytes, we treated control and PARP1-depleted terminally differentiated cardiomyocytes with the hypertrophic agonist Phenylephrine (PE) and assessed the expression of ANP as a marker of hypertrophic response. We observed an increase in ANP expression in PARP1-depleted cardiomyocytes compared to the controls. Importantly, there was a further increase upon PE treatment in PARP1-depleted cells, indicating that PARP1 depletion is associated with hypertrophic changes in cardiomyocytes **(Figure 5A)**. To verify this result, we performed inhibitor experiments with benzamide, a PARP1 inhibitor. Consistent with our previous findings, we observed that ANP expression was increased in the benzamide and PE-treated cardiomyocytes, indicating that PARP1 inhibition augments the development of a hypertrophic response in terminally differentiated cardiomyocytes (**Figures 5B, C**). Together, these results indicate that depletion of PARP1 in cardiomyocytes is associated with the development of hypertrophy in these cells.

**Figure 5.**
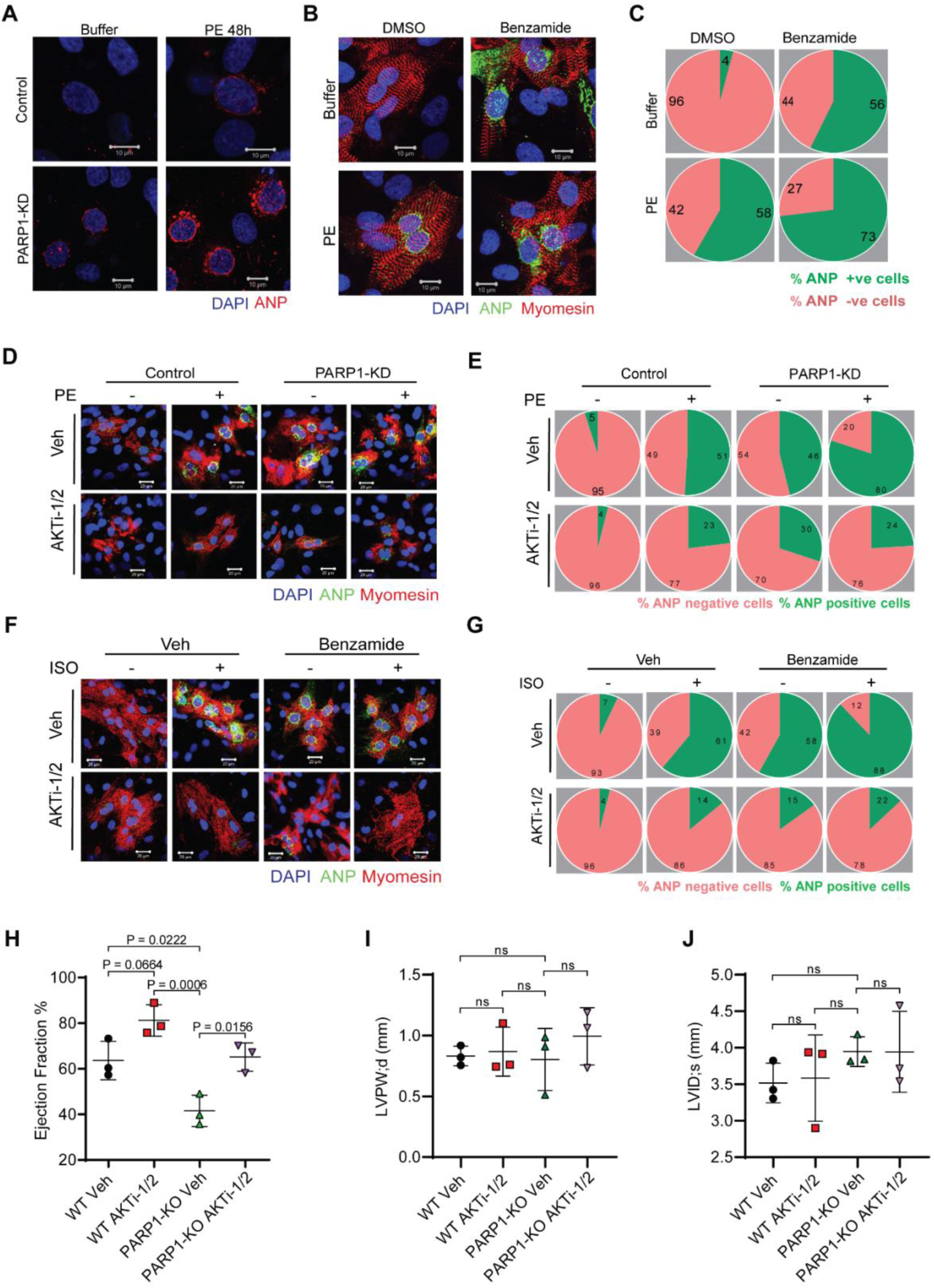
Akt inhibition improves cardiac function in PARP1-KO mice. **(A)** Representative confocal images showing ANP perinuclear localization in NRCM transfected with either Scrambled siRNA or PARP1-specific siRNA and further treated with Phenylephrine (PE) or Vehicle (Buffer) for 48 hours. The nucleus is stained with DAPI (blue), and ANP is stained in red. Scale bar = 10 µm (Scramble Buffer n=52, scramble PE n=78, PARP1KD Buffer n=36, PARP1 KD PE n=66). (**B)** Representative confocal images showing ANP perinuclear localization in NRCM Treated with either vehicle or PARP1 inhibitor (Benzamide) under basal or PE treatment for 48Hr. The nucleus is stained with Dapi (Blue), Myomesin (Red), and ANP (Green) Scale bar = 10 µm. **(C)** Representative graph showing the quantification of the percentage of ANP-positive and negative nuclei in control and PARP1-inhibited NRCM at the basal level and after 48 hours of PE treatment, as shown in Figure 3M. ANP-positive cells in the Pie chart are represented in green, and the percentage of ANP-negative cells is represented in red. N =3, (Veh Buffer *n* = 74, Veh PE *n* = 64, Benzamide Buffer *n* = 67, Benzamide PE *n* = 55). (**D)** Representative immunofluorescence image of neonatal rat cardiomyocytes (NRCMs) depicting ANP perinuclear localization under control or PARP1 depletion, upon treatment with Vehicle or Phenylephrine (PE), and further treatment with vehicle or Akt inhibitor (AKTi-1/2). Cells are stained with DAPI (Blue), ANP (Green), and Actinin (Red). Scale bar-20µm. **(E)** A representative graph showing the quantification of the percentage of ANP-positive and negative nuclei in control and PARP1-depleted NRCM at the basal level and after the PE treatment, followed by Vehicle (Veh) or AKTi-1/2 treatment, as shown in Figure 5D. ANP-positive cells in the Pie chart are represented in Green, and the percentage of ANP-negative cells is represented in Red. **N= 3,** (Veh Buffer veh n = 55,d Veh PE veh n = 71, PARP1 KD Buffer veh n = 56, PARP1 KD PE veh n =50, scr Buffer AKTi-1/2 n =50, scr PE AKTi-1/2 n =13, PARP1 KD Buffer AKTi1/2 n=10, PARP1 KD PE AKTi1/2 n=42. (scr = scrambled siRNA). **(F)** Representative immunofluorescence image of neonatal rat cardiomyocytes (NRCMs) depicting ANP perinuclear localization under control or PARP1 inhibition, upon treatment with Vehicle or Phenylephrine (PE), and further treatment with vehicle or Akt inhibitor (AKTi-1/2). Cells are stained with DAPI (Blue), ANP (Green), and Actinin (Red). Scale bar-20µm. **(G)** A representative graph showing the quantification of the percentage of ANP positive and negative nuclei in control and PARP1-inhibited NRCM at the basal level and after the PE treatment followed by Vehicle or AKTi-1/2 treatment, as shown in Figure 5F. ANP-positive cells in the Pie chart are represented in green, and the percentage of ANP-negative cells is represented in red. (Veh Buffer veh n = 58, Veh ISO veh n = 66, Benzamide Buffer veh n = 24, Benzamide ISO Veh n =41, Veh Buffer AKTi1/2 n=53, Veh ISO AKTi1/2 n=51, Benzamide Buffer AKTi1/2 n =71, Benzamide ISO AKTi1/2 n=23. (**H-J)** Scatter plot represents **(H)** Ejection fraction **(I)** Fractional shortening **(J)** Left Ventricular Internal Diameter diastole, LVID;d of WT and PARP1-KO post 2-week continuous injection of vehicle or Akt inhibitor (AKTi-1/2). n = 3 mice. Data information: In **(H-J)** data are represented as mean ± S.D.*P≤0.05; two-way ANOVA with Tukey’s multiple comparisons test was used to calculate p values. n = 3mice.

### Akt inhibition suppresses hypertrophic response in PARP1-deficient cardiomyocytes

To validate the role of PARP1-based regulation of Akt-mediated signalling and its implication in the hypertrophic heart, we treated PARP1-depleted terminally differentiated cardiomyocytes with the Akt inhibitor, AKTi-1/2. We hypothesized that if PARP1 regulates cardiac homeostasis via Akt, ANP expression would be reduced in the PE and AKTi-1/2 treated PARP1-knockdown cardiomyocytes. As expected, PE-treated PARP1-KD terminally differentiated cardiomyocytes showed higher ANP expression when compared to their respective controls. Moreover, we observed that ANP expression was reduced upon AKTi-1/2 treatment in PARP1-KD cells, even in the absence of PE, indicating that Akt inhibition in PARP1-depleted conditions alone is sufficient to partially rescue the hypertrophic response. Notably, the ANP expression was further reduced in PE-treated PARP1-KO cells, thereby verifying our findings **(Figures 5D, E)**. These results indicate that PARP1 regulates cardiomyocyte growth in an Akt-dependent manner. We further verified these results using another *in vitro* model, which involved benzamide treatment for PARP1 inhibition and isoproterenol (ISO) treatment for induction of a hypertrophic response in terminally differentiated primary cardiomyocytes. Similar to our previous findings, both untreated and ISO-treated PARP1-KD cardiomyocytes showed higher ANP expression compared to their respective controls. Notably, we observed that ANP expression was reduced in AKTi-1/2 and benzamide-treated cells compared to benzamide-treated cells, even in the absence of ISO, indicating that Akt inhibition in PARP1-inhibited conditions is sufficient to partially rescue the hypertrophic response **(Figures 5F, G)**. These results suggest that Akt inhibition rescues the hypertrophic response induced upon PARP1 depletion in terminally differentiated primary cardiomyocytes. Together, our *in vitro* models suggest that PARP1 depletion may induce cardiac hypertrophy by modulating Akt signalling.

### Akt inhibition improves contractile function in PARP1-deficient mice

We investigated whether inhibiting Akt could rescue the cardiac remodeling and failure observed in PAPR1-KO mice. We injected wild-type and PARP1-KO mice with the Akt inhibitor, AKTi-1/2. Following this, we analysed cardiac structure and function in these mice via echocardiography. Although we found no improvement in cardiac structural parameters, such as internal diameter and wall thickness, we observed that the cardiac dysfunction in PARP1-KO mice was rescued upon treatment with AKTi-1/2, as indicated by restored cardiac ejection fraction **(Figures 5H-J).** This suggests that cardiac dysfunction observed in PARP1-KO mice can be rescued upon Akt inhibition, suggesting that PARP1 regulates cardiac function, at least partially, in an Akt-dependent manner.

## Discussion

Our results suggest a significant role for PARP1 in regulating Akt signalling in the mouse heart, which protects mice against aging-associated heart failure. Our findings reveal two distinct mechanisms by which PARP1 regulates Akt signalling, involving the direct interaction of PARP1 with Akt and the transcriptional regulation of PTEN expression by PARP1. We believe that these two distinct mechanisms provide rigorous control over Akt signalling, as this signalling pathway plays a critical role in cardiac growth and aging.

Although many studies have examined the involvement of PARP1 in cardiac function under stress [42–44], its contribution to age-related cardiomyopathy has remained largely unresolved. Experimental studies indicate that PARP1 contributes significantly to cardiac dysfunction associated with diabetes [44]. Diabetic mice exhibit increased PARP1 expression and activity in the heart, indicating its activation in response to metabolic and oxidative stress. Blocking PARP1 activity reduces high–fat–diet–induced susceptibility to ischemic damage, resulting in significant reductions in cardiac inflammation and fibrosis [44]. Moreover, Olaparib-mediated suppression of PARylation reduces H₂O₂-driven oxidative injury and cell death in cardiomyocytes, while improving post-transplant cardiac performance in rats [42]. Contrary to these earlier findings suggesting a detrimental influence of PARP1 on cardiac health, our findings indicate that PARP1 plays a protective role in age-associated cardiac decline. It is quite possible that PARP1 might function differently in acute and chronic stress conditions. Acute cardiac stress induced by oxidative and genotoxic stress, as well as ischemia and reperfusion injury, may hyperactivate PARP1, leading to depletion of cellular NAD and induction of apoptosis [43, 45]. In this situation, inhibition of PARP1 may preserve NAD and protect cardiomyocytes from cell death, thereby offering cardio-protection [43, 45–48]. However, chronic inhibition of PARP1 may activate pro-aging signalling pathways, including the Akt kinase, leading to the development of aging-related cardiac hypertrophy and failure. Thus, PARP1 plays a critical role in maintaining cardiac homeostasis by balancing cell growth and death during stress conditions.

PARP1 has been the focus of extensive research because of its central roles in DNA repair, transcriptional control, maintenance of genomic integrity, and regulation of cell death pathways [8, 49–51]. However, the involvement of PARP1 in specific cellular signalling pathways remains poorly understood. Previous studies have emphasized the role of PARP1 in regulating MAPK signalling [52]. Additionally, PARP1 plays a crucial role in regulating the deacetylase activity of SIRT1 [53]. Moreover, evidence from various reports also implicates PARP1 in transcriptional regulation within of NF-κB pathway [52]. In the current study, our findings establish a direct connection between PARP1 and Akt signalling, an evolutionarily conserved pathway that responds to nutrients and is involved in aging [54–57]. Surprisingly, all these molecules are linked to the aging process [58–64], suggesting a key role for PARP1 in regulating various aging-related signalling pathways involved in cardiac aging. It is not surprising that PARP1-KO mice develop accelerated aging and a reduced lifespan [10].

Dysfunction of Akt contributes to serious health conditions such as cardiovascular disease, cancer, diabetes, and neurological disorders [65]. Our results indicate that PARP1 deficiency leads to sustained activation of Akt signalling, a key contributor to aging and age-related diseases [66]. Studies in cardiac-specific inducible Akt models suggest that short-term Akt1 upregulation supports physiological hypertrophy, whereas long-term activation culminates in hypertrophy with pathological consequences, notably reduced contractile performance [67]. Additionally, Akt appears to exacerbate aging-induced defects in cardiac structure and function by disrupting autophagy [68]. Thus, it is evident that chronic and sustained activation of Akt due to PARP1 deficiency could induce aging-related cardiac dysfunction.

While this study has several strengths, including the identification of the role of PARP1 in cardiac aging and the characterization of Akt residues regulated by PARP, it also has some limitations. This study utilized whole-body PARP1-KO mice, rather than cardiomyocyte-specific knockout mice. Also, this study has not confirmed the Akt residues that are PARylated by PARP1 using mass spectroscopy. This study also did not fully uncover the molecular mechanisms involved in PARP1-mediated regulation of phosphatases. Future studies are needed to elucidate the role of PARP1 in the aging of major cell types, including cardiomyocytes, fibroblasts, cardiac stem cells, and endothelial cells. Nevertheless, our study reveals a previously unrecognized role for PARP1 in regulating cardiac aging, underscoring the need for a targeted approach to utilizing PARP1 inhibitors in the treatment of cardiovascular disorders in humans.

## Methodology

### Animal experiments

All animal experiments were performed with the approval of the Institute Committee for Control and Supervision of Experiments on Animals (CPCSEA), Government of India, and in accordance with the norms of the Institutional Animal Ethics Committee (IAEC) of the Indian Institute of Science (CAF/Ethics/818/2021). Individually ventilated cages were used to maintain the mice, and a standard chow diet was fed to them under a 12-hour light/dark cycle. Whole-body PARP1 knockout mice were obtained from Jackson Laboratory, USA (Jackson Laboratory, Cat# 002779). For genotyping, whole-body PARP1 knockout mice, the PCR cycling conditions were as follows: initial denaturation at 95°C for 3 minutes; 40 cycles of 94°C for 30 seconds, 56°C for 45 seconds, and 72°C for 45 seconds; followed by a final extension at 72°C for 5 minutes.

### Echocardiography

Transthoracic echocardiography (TTE) was performed non-invasively by a trained operator using a high-resolution ultrasound system (Vevo 1100 Imaging System, FUJIFILM VisualSonics), in accordance with ASE guidelines for small animals. Mice were anaesthetized with 1.5% to 2.0% isoflurane, maintaining physiological stability by monitoring body temperature at 37.0±0.5°C and heart rate between 400 and 500 beats per minute. Image acquisition was performed with two-dimensional (2D) guidance in the Parasternal Long-Axis (PLAX) and Parasternal Short-Axis (PSAX) views. These images were used to calculate systolic function parameters, specifically Left Ventricular Fractional Shortening (LVFS) and Ejection Fraction (LVEF), as well as estimated LV Mass. All parameters were averaged over at least three stable cardiac cycles for offline analysis.

### AKT inhibitor experiments

9- to 10-month-old cohorts of wild-type and PARP1 knockout mice were injected with the vehicle and the AKTi 1/2 inhibitor (dose: 1mg/kg/day) intraperitoneally for a duration of 14 days. The echocardiographic parameters were assessed at the end of the regimen, as described in the previous section.

### Primary cardiomyocyte culture

Neonatal Rat Primary cardiomyocytes (NRCMs) were derived from neonatal rat pups and cultured as described in our previous publication [69]. Briefly, the pups were anaesthetized using isoflurane and then sacrificed. The hearts were harvested into PBS-glucose. The collected heart tissue was minced and to digested using collagenase II and 0.2% trypsin. At the end of each digestion cycle, the cell pool was collected in horse serum. Upon complete digestion of heart tissue, the cell pool was pre-plated for 45 minutes in a 100mm dish to separate the fibroblasts from the cardiomyocytes. After the incubation period, the unattached cells (cardiomyocytes) were harvested from the dishes, centrifuged, and then seeded into 0.2% gelatin-coated well plates. For the terminal differentiation of NRCMs we utilized a previously published protocol [70]. Briefly, cells were treated with a defined low-glucose (Gibco™ Cat#11885084), serum-free (LGSF) medium instead of conventional high-glucose, serum-rich conditions; additionally, cells were supplemented with a free fatty acid (FFA)-albumin complex (Sigma Aldrich, Cat#L9655) to enhance cardiomyocyte maturation [71]. Cells were cultured in these conditions up to 5 days to obtain terminally differentiated cardiomyocytes.

### Cell culture

HEK293 and HeLa cells were procured from ATCC and maintained in Dulbecco’s modified Eagle’s medium complemented with 10% fetal bovine serum (FBS) and 1X antibiotic/antimycotic solution, at 37°C with 5% carbon dioxide. Cells were seeded overnight to reach 70-80% confluency at the time of transfection. PARP1 knockdown was performed by transfecting cells with PARP1-specific siRNA using Lipofectamine 3000 (Thermo Fisher Scientific) according to the manufacturer’s instructions. Cells were incubated for 48 hours post-transfection, and knockdown efficiency was confirmed by qRT-PCR and Western blot analysis. PARP1 overexpression was performed by transducing cells with hPARP1 adenovirus for a period of 12 hours. Western blotting and qPCR were performed to experimentally validate the overexpression of PARP1.

### RNA isolation and qPCR analysis

Cultured cells and mouse heart tissue were homogenised, and RNA was isolated using TAKARA RNAiso Plus (Cat#9108). 500 ng of the total isolated RNA was used for cDNA synthesis, performed using the Bio-Rad iScript cDNA synthesis kit (Cat#1708890) as per the manufacturer’s protocol, and qPCR was performed using SYBR-Green PCR master mix in the QuantStudio^TM^ Real-Time PCR system. Sequences of primers used in this study are provided in **Table 1**.

**Table 1.**
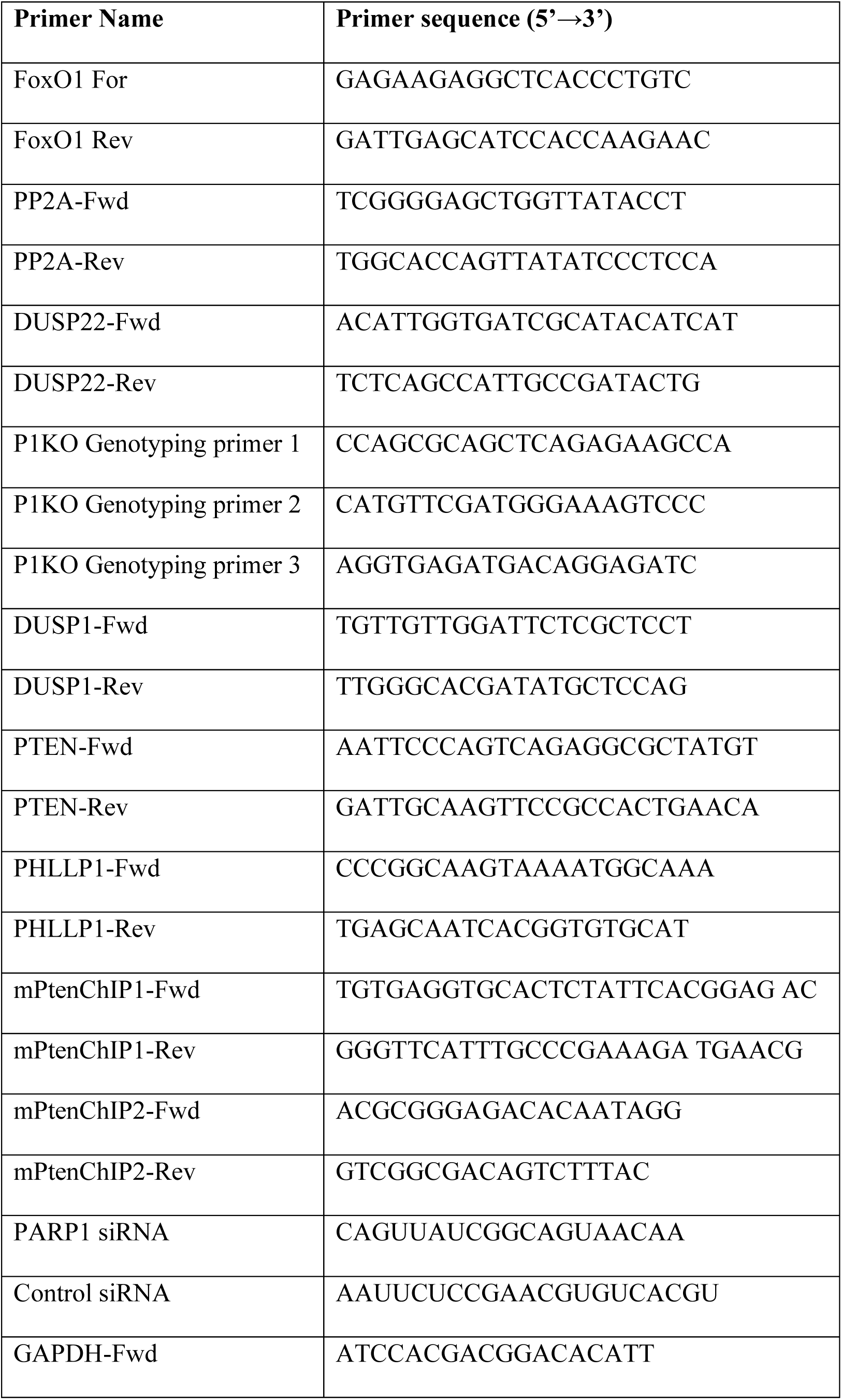
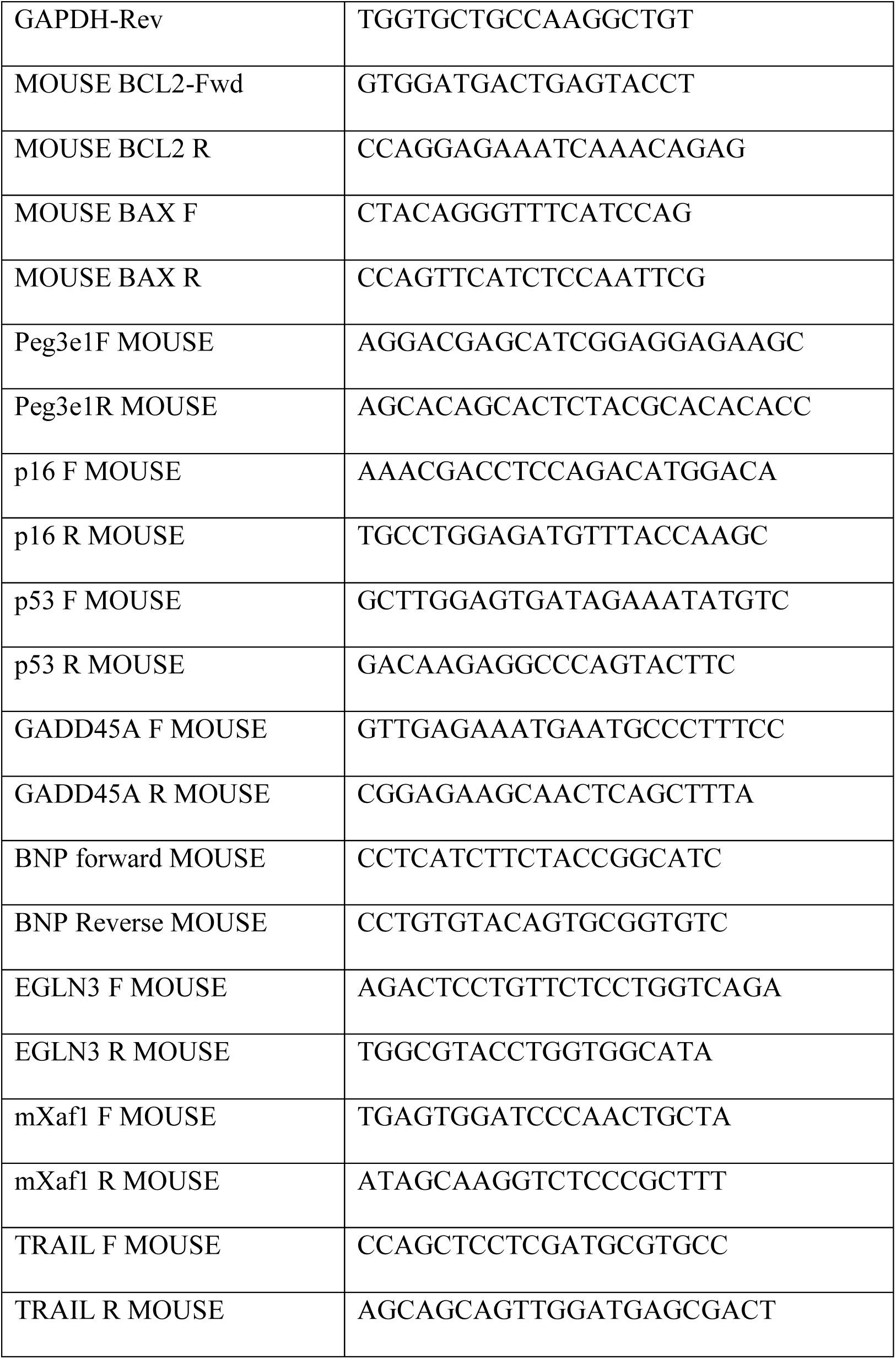

### RNA sequencing

RNA was isolated from the heart tissue of whole-body PARP1 knockout mice as described in the previous section. The concentration and quality of the isolated RNA were evaluated using a NanoDrop Spectrophotometer (ThermoFischer), and high-quality RNA samples were sent for sequencing to MolSys Pvt. Ltd. Raw FASTQ files for wild-type (WT) and knockout (KO), RNA-seq samples were subjected to sequencing quality assessment using *FastQC* (v0.12.1; http://www.bioinformatics.babraham.ac.uk/projects/fastqc/). The latest mouse reference genome assembly (GRCm39) and corresponding gene annotation file in GTF format were obtained from the GENCODE project (Release M36; https://www.gencodegenes.org/). Reads were aligned to the mouse reference genome (GRCm39) using *STAR* (v2.7.11)[72]. BAM files of technical replicates were merged with *SAMtools* (v1.18)[73]. *featureCounts* (v2.0.6) was used to perform gene-level quantification[74]. Differential gene expression analysis was conducted using *DESeq2* (v1.43.5)[75]. Raw read counts were normalized using estimated size factors, and differential expression was assessed using a negative binomial generalized linear model that incorporated shrinkage estimation of dispersion and fold changes. Genes were considered significantly differentially expressed at │log_2_FC│ ≥ 0.58 and *p-*value < 0.05 calculated using Wald’s test. Principal component analysis (PCA) was used to evaluate the variability among biological replicates. Variance-stabilized transformed (VST) gene expression data from *DESeq2* were used for heatmap visualization with the *ComplexHeatmap* R package (v2.22.0)[76]. Volcano plots depicting log_2_ fold changes of genes were generated using the *ggplot2* R package (v3.5.1)[77]. *GSEA* (v4.3.3) was used to perform gene set enrichment analysis (GSEA)[78], using ranked gene lists from the differential expression analysis to assess enrichment of biological pathways against the Molecular Signatures Database (MSigDB).

### Immunoblotting

Frozen mouse tissues and cell cultures were homogenised and lysed using RIPA lysis buffer (1 mM EDTA, 2.5 mM sodium pyrophosphate, 150 mM NaCl, 20 mM Tris–HCl, pH 7.5, 1mM PMSF, 1mM sodium orthovanadate, 1mM EGTA, 1% NP-40, 1X protease inhibitor cocktail, and 1% sodium deoxycholate). The lysate was vortexed for 30 minutes at 4°C and then centrifuged for 15 minutes at 12,000 rpm, 4°C. Protein concentration estimation was performed using Bradford reagent (BIO-RAD, Cat#5000205). Samples were prepared accordingly and then heated at 95°C for 5 minutes in 2X Laemmli buffer supplemented with 5% β-mercaptoethanol for denaturation. Proteins were separated by 12% SDS-PAGE and subsequently transferred to 0.45 mm PVDF membranes via overnight wet transfer (20V). Membranes were blocked with 5% skimmed milk in TBST. Target proteins were detected using the following primary antibodies: PAR (Calbiochem, Cat#AM80), Akt (SantaCruz, Cat #sc-8312, Cell Signalling Technology Cat#4691), Phospho-Akt (Cell Signalling Technology Cat#Ser473-4060), Phospho-FoxO1 (Cell Signalling Technology Cat#Ser256-9461), FKHR (SantaCruz, Cat#sc-11350), PARP1 (SantaCruz, Cat#sc-7150), GAPDH (SantaCruz, Cat#sc-25778), GSK-3β (Cell Signalling Technology Cat#9315), Phospho-GSK-3β (Ser9) (Cell Signalling Technology Cat#9336), β-Actin-HRP Conjugate (Cell Signalling Technology Cat#12262). Signal was visualised using a chemiluminescence substrate (Thermofischer Cat#34080) on a Bio-Rad ChemiDoc Touch system. Protein expression analysis and image processing were carried out using Image Lab (Bio-Rad) software.

### Confocal Microscopy

Samples were prepared for confocal microscopy as described previously [79]. Briefly, gelatin-coated coverslips were seeded with 0.2 million primary cardiomyocytes or HeLa cells in 24-well plates. For subsequent staining for immunofluorescence, the media from the cells were removed, washed with ice-cold PBS, and fixed using 4% paraformaldehyde for 15 minutes at room temperature. Blots were blocked using 5% BSA at room temperature for 1 hour to avoid nonspecific binding. The blocked cells were then incubated with the required primary antibody at 4°C. After incubating the cells with the primary antibody overnight (12-16 hours), the cells were washed three times with PBST and then incubated with the specific fluorophore-conjugated secondary antibody for one hour at room temperature. After incubation with the secondary antibody, the cells were washed three times with PBST and then mounted using Prolong Gold antifade reagent along with DAPI/Hoechst (Molecular Probes). A Zeiss LSM 880 with Airyscan or a Zeiss LSM 710 microscope was used to capture the images. After capturing the fluorescence images, they were processed using ZEN Black (Carl Zeiss) and ImarisViewer 10.0.1 software. Subsequent image quantification was performed using ImageJ Fiji (developed by the National Institutes of Health, USA).

### Chromatin immunoprecipitation assay

The protocol for *in vivo* Chromatin immunoprecipitation in this study was adapted from our previously published work [80]. For the assay, whole-heart tissue harvested from wild-type mice was homogenized using an electric homogenizer in 1X PBS at room temperature. Crosslinking of the homogenized tissue was performed in a 1% formaldehyde solution with continuous shaking for 15 minutes. The crosslinked product was then quenched using glycine to achieve a final concentration of 150 mM. The quenched tissue was centrifuged at 2000g at 4°C for 10 minutes. The cell pellet was then lysed using the cell and nuclear lysis buffer. The lysed product was then sonicated with the standardised on/off cycle of 25s (on) and 40s (off) for 3-4 cycles. The size of the sheared product was determined by running it on an agarose gel, and the size ranged from 200 bp to 600 bp. 50μg of the chromatin was immunoprecipitated using a PARP1 primary antibody. Immunoprecipitated chromatin was then de-crosslinked and eluted using ChIP elution buffer at 50°C with vortexing for 30 minutes at 700 rpm. The eluted DNA was then used for qPCR-based analysis of the PTEN primer.

### Histology Analysis

Heart tissue was fixed for 72 hours in 10% neutral buffered formalin [81]. The fixed tissue was processed, embedded in paraffin, and sectioned using the microtome to obtain sections of 5 μm thickness. Cell size was measured using wheat germ agglutinin (WGA) stain by standard techniques, and images were acquired with confocal microscopy, followed by quantification using NIH ImageJ software. Fibrosis was detected using Masson’s trichrome staining and quantified manually through visual assessment of the fibrosis.

### Co-immunoprecipitation assay

293 cells were washed with ice-cold PBS and harvested in ice-cold RIPA buffer (50 mM Tris-Cl, 150 mM NaCl, 1% NP-40, 0.25 % sodium deoxycholate, 0.1 % SDS, 1 mM EDTA,1mM Na3VO_4_, 2.5 mM sodium pyrophosphate, protease inhibitor cocktail (Roche, Cat#11 697 498 001), 1 mM PMSF, 1 µM ADP-HDP) and lysed by vigorous vortexing for 30 min at 5 min interval each which was followed by centrifugation for 15 min at 13000 rpm. The supernatant was collected, and protein concentration was quantified using Bradford reagent (Bio-Rad, Cat # 50006) at 595 nm. 500 µg of total protein was incubated overnight at 5 rpm, 4°C with 2 µg of antibodies: PAR antibody (Calbiochem, Cat#AM80) and AKT antibody (sc-1618). 20 µl of protein A bead (Pierce, Cat#20334) was washed three times with RIPA buffer for 30 sec at 1000 rpm. A whole-cell lysate that was incubated overnight with the antibody was added to the washed protein A beads and kept on rotation at 5 rpm, 4°C for 2 hours. Beads were washed three times with ice-cold PBS, and bead-bound proteins were eluted by adding 2x Laemmli sample buffer (Bio-Rad, Cat#161-0737) and boiled at 96 °C for 5 min. The eluted proteins were analysed after immunoblotting using Akt (sc-8312), PARP1 (sc-7150), and GAPDH (sc-25778) antibodies.

### T7 pull-down assay

293 cells were transfected with empty vector or T7 Akt or T7 Akt mutants, T7 AKT E17A, T7 Akt E40A or T7 Akt E49A expression plasmids using Lipofectamine 2000 (Invitrogen). 24 hours post-transfection, cells were serum-starved for 18 hours and then induced for 15 minutes with 10% fetal bovine serum (FBS). Cells were harvested in ice-cold RIPA buffer as described above. Agarose beads conjugated to T7 antibody (Novagen, Cat#69026) was washed 3 times with RIPA and incubated with 200 µg of total protein lysate for 2-3 hr at 4°C at 5 rpm. Beads were washed 3 times with PBS, and protein was eluted from beads by adding 2x Laemmli sample buffer and boiling for 5 min at 96°C, followed by immunoblotting.

### Site Directed Mutagenesis

We used QuikChange II Site-Directed Mutagenesis Kit (Agilent Technologies) and performed SDM in T7-tagged Akt1 and GFP Akt PH plasmids to generate T7-Akt1 E17A, T7-Akt1 E40A, and T7-Akt1 E49A, GFP Akt1 E17A, GFP Akt1 E40A, and GFP Akt1 E49A mutants according to the manufacturer’s protocol. These mutants contain a single point mutation of the indicated glutamine (E) residues, where Glutamine has been replaced by Alanine. The primers used to generate SDMs are listed below.

E17A For: CCAGGTCTTGATGTACGCCCCTCGTTTGTGCAG

E17A Rev: CTGCACAAACGAGGGGCGTACATCAAGACCTGG

E40A For: CATCCTGCGGCCGCGCCTTGTAGCCAATG

E40A Rev: CATTGGCTACAAGGCGCGGCCGCAGGATG

E49A For: GTTGAGGGGAGCCGCACGTTGGTCCAC

E49A Rev: GTGGACCAACGTGCGGCTCCCCTCAAC.

### Statistical analysis

The data were reported as mean ± standard deviation. Datasets were subjected to Kolmogorov-Smirnov and Shapiro-Wilk tests to determine if they were normally distributed. For normally distributed datasets, statistical analysis was performed using the Student’s t-test (for two groups) or ANOVA (for three or more groups), followed by Tukey’s or Sidak’s multiple comparison tests. Mann–Whitney U or Kruskal–Wallis tests, followed by Dunn’s comparisons, were used for non-normally distributed data. A p-value of ≤0.05 was considered statistically significant. Graph Pad Prism 8.4.2. was used for data analysis and graph plotting.

## RESOURCE AVAILABILITY

### Lead Contact

Further information and requests for resources and reagents should be directed to the Corresponding Author, N. Ravi Sundaresan.

### Materials availability

All unique/stable reagents generated in this study are available with the Lead Contact (N. Ravi Sundaresan) with a completed Materials Transfer Agreement.

### Data and code availability

This study generated the unique datasets/code SAMN52643000, SAMN52643001, SAMN52643002, SAMN52643003, SAMN52643004, SAMN52643005. The source data for figures in the paper are available as **Supplemental file 1.**

## ACKNOWLEDGEMENTS

The Central Animal Facility, the Microtome Facility, Division of Biological Sciences, and the Confocal Facility, Division of Biological Sciences, Department of Microbiology and Cell Biology at the Indian Institute of Science, Bengaluru; and the Mouse Genome Engineering Facility at the National Centre for Biological Sciences, India, are acknowledged for their services and the technical help.

## AUTHOR CONTRIBUTIONS

NRS conceived and designed the study and prepared the final draft. SM and TSS designed the experiments. SM and TSS analysed the experiments. AJS, TSS, AM, and AYB wrote the manuscript. TSS and SM performed the primary cardiomyocyte culture, confocal microscopy, RNA isolation, RT-PCR, and quantification of the results. TSS and SM bred the PARP1 knockout mouse line. TSS performed genotyping of the PARP1 knockout mouse line. TSS prepared samples for RNA sequencing analysis. TSS performed the qPCR. SM, AJS, and DN performed immunoblotting. TSS performed histological experiments. AC performed the RNA-sequencing data analysis. TSS and SVR performed co-immunoprecipitation experiments. All authors have cross-verified the original data in the manuscript.

